# A Systematic Approach for Explaining Time and Frequency Features Extracted by CNNs from Raw EEG Data

**DOI:** 10.1101/2022.02.08.479555

**Authors:** Charles A. Ellis, Robyn L. Miller, Vince D. Calhoun

## Abstract

In recent years, the use of convolutional neural networks (CNNs) for raw electroencephalography (EEG) analysis has grown increasingly common. However, relative to earlier machine learning and deep learning methods with manually extracted features, CNNs for raw EEG analysis present unique problems for explainability. As such, a growing group of methods have been developed that provide insight into the spectral features learned by CNNs. However, spectral power is not the only important form of information within EEG, and the capacity to understand the roles of specific multispectral waveforms identified by CNNs could be very helpful. In this study, we present a novel model visualization-based approach that adapts the traditional CNN architecture to increase interpretability and combines that inherent interpretability with a systematic evaluation of the model via a series of novel explainability methods. Our approach evaluates the importance of spectrally distinct first-layer clusters of filters before examining the contributions of identified waveforms and spectra to cluster importance. We evaluate our approach within the context of automated sleep stage classification and find that, for the most part, our explainability results are highly consistent with clinical guidelines. Our approach is the first to systematically evaluate both waveform and spectral feature importance in CNNs trained on EEG data.

## Introduction

In recent years, the use of convolutional neural networks (CNNs) in the analysis of raw electroencephalography (EEG) data has grown considerably. These classifiers have the advantage over standard machine learning and deep learning classifiers paired with manual feature extraction in that they don’t require any prior assumptions about the important features within the data and that they automate feature extraction. While this is the case, pairing automated feature extraction with raw time-series data also causes problems with explainability, which is highly important in sensitive domains like healthcare. As such, novel methods for explaining CNNs trained on raw EEG data are needed. In this study, we present a novel approach that pairs a CNN architecture adapted for increased interpretability with a series of systematic model perturbations that provide valuable insight into the features extracted by the CNN and the relative importance of those features. Unlike previous approaches that have mainly provided insight into key frequency feature extracted by CNNs, we provide insight into both the frequencies and waveforms extracted by CNNs.

Prior to the recent trend of applying CNNs to raw electrophysiology data for automated feature extraction, it was common to manually create features and apply machine learning or deep learning approaches with traditional explainability methods when analyzing electrophysiology data. The user-created features typically reflected either time domain or frequency domain (1–5) aspects of the data. A strength of this approach was that it could be applied alongside explainability methods initially developed outside the domain of electrophysiology analysis like layer-wise relevance propagation (LRP) (6), Grad-CAM (7), and activation maximization (8). However, the use of user-selected input features also inherently limited the available feature space, thereby limiting the potential performance of classifiers.

As such, more studies have begun to apply CNNs to raw electrophysiology analysis. While this application can improve model performance, applying traditional explainability methods to raw time-series samples makes it very difficult to know what time or frequency features are extracted by classifiers and to draw global conclusions about the importance of extracted features (9). It should be noted that this difficulty is not applicable to identifying spatial importance (10) or modality importance (11–14) in multichannel or multimodal classification, respectively. However, this difficulty is applicable when trying to understand the temporal and spectral features extracted by classifiers.

In response to the need for improved explainability in CNNS applied to raw electrophysiology data, a new field of explainability for CNN-based raw electrophysiology classification has developed. The vast majority of these methods provide insight into frequency-based features extracted by CNNs. These methods can loosely be divided into 4 categories: (1) interpretable architectures, (2) activation maximization approaches, (3) perturbation approaches, and (4) model visualization approaches. Interpretable architectures involve structuring filters in the first convolutional layer such that they only extract spectral features (15,16). While these methods are very innovative, they still inherently restrict the feature space to frequency features. Several studies have presented methods that use activation maximization to identify spectral features that maximize activation of the CNN (17–19). Two studies examined the effect of sinusoids at different frequencies upon activations of nodes in the early layers of the classifier (17)(19), and the remaining study varied the spectral representation of a sample until it maximized the activation of the final output node (18). Other existing studies involve perturbing canonical frequency bands of samples and examining the effect upon the predictions (20)(21) or performance of a classifier (22). The last category of methods involves training a CNN with long first-layer filters that can be converted to the frequency domain after training and visualized to examine the spectral features extracted by the model (19). While these methods do provide useful insight into extracted spectra, they do not provide effective insight into extracted time-domain features.

For the most part, existing approaches that provide insight into insight into time-domain features of EEG are of limited utility. One study perturbs windows of a time-domain sample (17). However, when datasets consist of thousands of samples and there is no way to combine insights from the perturbation of each sample, that approach does not provide useful global conclusions on the nature of the time-domain features extracted. Another study used activation maximization to optimize the spectral content of a sample (18). While the method does yield a sample in the time domain that maximizes activation for a particular class, it does not provide insight into the relative importance of different time domain features. A couple of other studies have used activation maximization for insight into the time domain (23,24).

However, they were only applied to networks trained on samples that were around 30 time points long, and EEG samples can be hundreds to thousands of time points long. Additionally, a previous study showed that these methods do not generalize well to sample lengths relevant for EEG analysis (18). It should be noted that existing explainability methods can be applied to some forms of electrophysiology like electrocardiograms (ECG) that have regularly repeated waveforms (25,26). However, these applications rely upon the regular repetition of waveforms, and that repetition is not present in forms of electrophysiology like resting-state magnetoencephalography and EEG. A method which provides useful insight into time domain features (27) extracted by CNNs for EEG involves training a CNN with filters in the first layer that are long enough to extract distinct waveforms (19). The filters can then be visualized and perturbed to examine the relative importance of each waveform within the filters to the classifier performance. We developed this method in a previous study (27), and we expand upon it here to provide a systematic approach for explaining CNNs trained on raw electrophysiology time-series.

In this study, we use sleep stage classification as a testbed to demonstrate the utility of our approach. Sleep stage classification has several noteworthy characteristics that make it ideal for our application. (1) The domain of sleep stage classification has well-characterized spectral and temporal features, so we can evaluate whether our explainability results are consistent with established scientific knowledge (28). (2) There are multiple large publicly available datasets within the domain of sleep stage classification that help with reproducibility of analyses (29–31). (3) Multiple studies have already presented explainability methods for the domain of sleep stage classification, which will make it easier to compare our findings with results from previous studies (17–20,32).

In summary, in this study, we present a novel systematic approach for gaining insight into the features extracted by CNNS from raw EEG data and into the relative importance of those features. We train a CNN for automated sleep stage classification with a publicly available dataset and structure the first layer of the architecture such that we can visualize the waveforms extracted by the model. We convert the first layer filters of the model to the frequency domain and identify clusters of filters with distinct spectral characteristics. We then examine the relative importance of each cluster of filters. After identifying the importance of each cluster, we examine how much of that importance is attributable to spectra and to multispectral waveforms within each cluster. Unlike previous methods that only provided insight into key spectral features, we provide insight into both key spectral features and waveforms. Figure 1 shows an overview of our methods.

**Figure 1.**
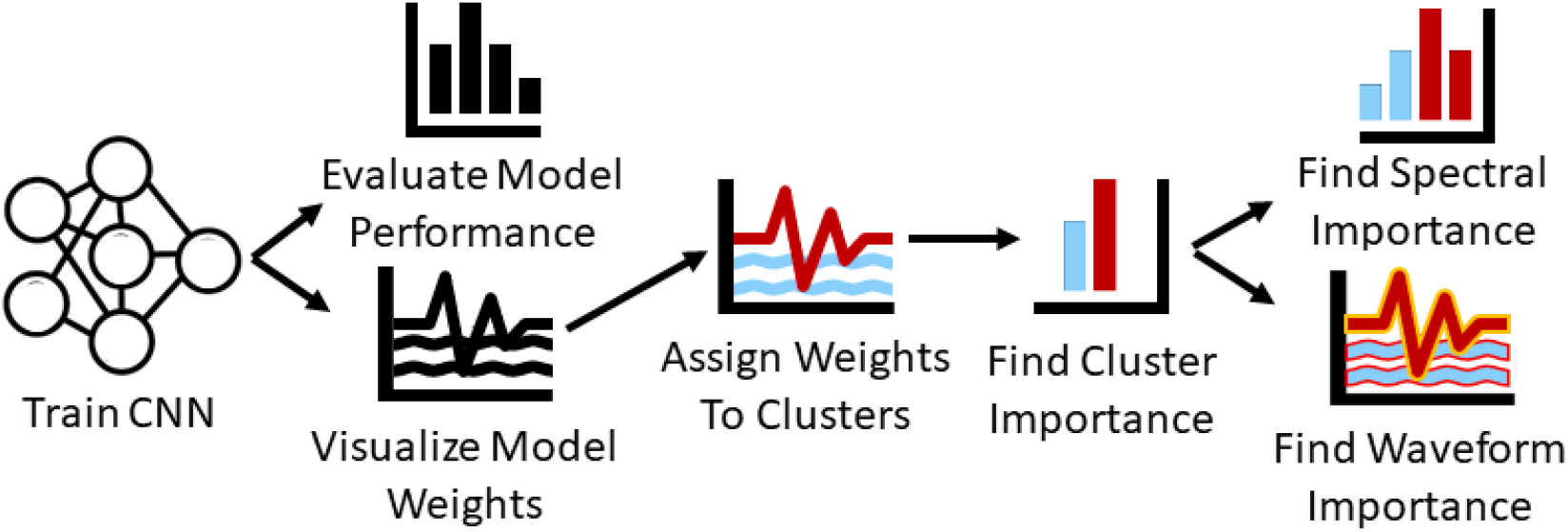
Overview of Methods. We train a CNN for sleep stage classification and evaluate its performance. We then visualize the first layer filters of the CNN, convert them to the frequency domain, and cluster them. After clustering the filters, we ablate each cluster to determine its importance and perturb their frequency and time domain representations to determine how much of cluster importance is attributable to the spectra and waveforms within each cluster.

## Methods

In this study, we use EEG sleep stage data to train a CNN. We then visualize the first layer filters of the CNN, convert them to the frequency domain, and cluster them. After clustering the filters, we ablate each cluster to determine their importance and perturb their frequency and time domain representations for insight into the importance of the spectra and waveforms within each cluster.

### Dataset and Data Preprocessing

In this study, we used the Sleep Cassette subset of the Physionet (33) Sleep-EDF Expanded Dataset (29). The Sleep Cassette subset contains 153 20-hour recordings from 78 healthy individuals. Each individual had 2 subsequent recordings of day-night periods while at home. The dataset includes electroencephalogram (EEG), electrooculogram (EOG), electromyogram (EMG), oro-nasal airflow, and rectal body temperature. However, in our study, we just used EEG from the FPz-Cz electrode recorded at 100 Hertz (Hz). The data was assigned by experts to Awake, REM, NREM1, NREM2, NREM3, and NREM4 stages in 30-second intervals.

We segmented the data into 30-second samples based on the expert assigned intervals. To alleviate data imbalances, we removed Awake data from the start of the recordings and part of the end of the recordings. Using clinical guidelines, we made NREM3 and NREM4 a single class. After removing samples, we separately z-scored the EEG data from each recording. Table 1 shows the resulting distribution of samples in each class.

**Table 1.** Distribution of Samples

|  | **Awake** | **NREM1** | **NREM2** | **NREM3** | **REM** | **Total** |
| --- | --- | --- | --- | --- | --- | --- |
| **Number** | 85,034 | 21,522 | 69,132 | 13,039 | 25,835 | 214,562 |
| **Percent** | 39.63 | 10.03 | 32.22 | 06.08 | 12.04 | 100 |

### Model Development

Here we discuss our model development and evaluation approach. We implemented our model architecture and training in Keras (34) and Tensorflow (35).

#### Architecture

We utilized a 1D-CNN architecture that had long filters in the first convolutional layer to make it easier to apply explainability methods and gain insight into extracted waveforms and frequency bands. The architecture that we used was originally developed in (19), and we adapted it to make it compatible with our shorter 30-second segments. Figure 2 shows our classifier architecture.

**Figure 2.**
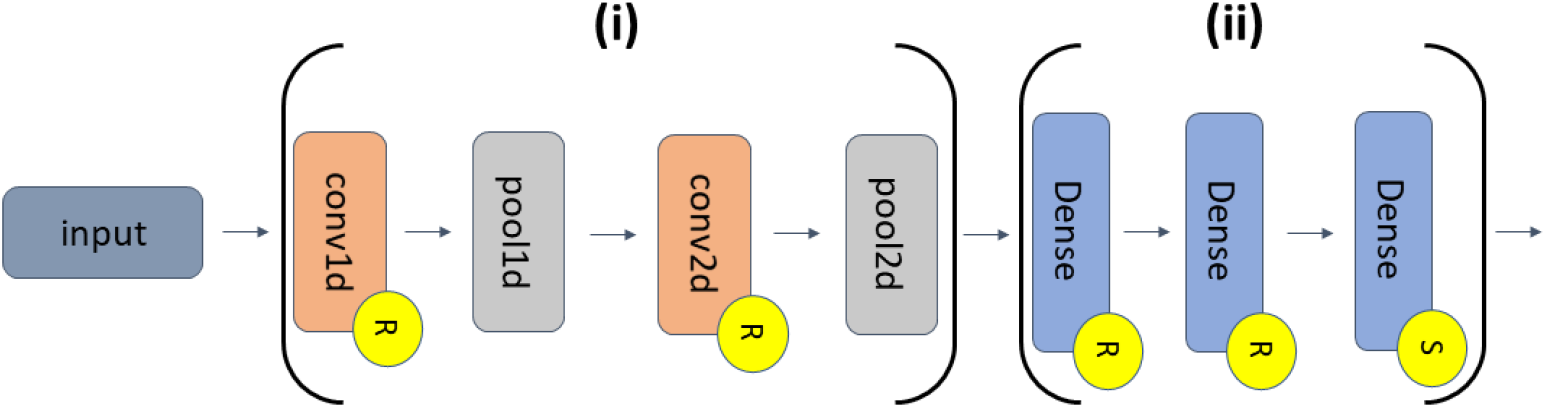
CNN Architecture. The feature extraction and classifier segments of the classifier are indicated by Sections (i) and (ii), respectively. (i) has of a 1D convolutional (conv1d) layer (30 filters, length of 200, stride of 1), a 1D max pooling (pool1d) layer (pooling size of 15, stride of 10), a conv2d layer (400 filters, 30 x 25 filter size, stride of 1 x 1), and a pool2d layer (pooling size of 10 x 1, stride of 1 x 2). (ii) has 2 dense layers (500 nodes) and an output dense layer (5 nodes). Layers with an “R” or an “S” have ReLU or softmax activation functions, respectively.

#### Cross-Validation and Training Approach

When developing our architecture and training our classifier, we used a 10-fold cross-validation approach. In each fold, we randomly assigned 63, 7, and 8 subjects to training, validation, and test groups, respectively. To address class imbalances, we weighted our categorical cross entropy loss function. We also used a stochastic gradient descent optimizer with a batch size of 100 and an adaptive learning rate with an initial value of 0.015 that decreased by 10% after each set of 5 epochs that did not have a corresponding increase in validation accuracy. We also applied early stopping if 20 epochs occurred without an increase in validation accuracy. The maximum number of training epochs was 30. Additionally, we shuffled the training data between each epoch.

#### Performance Evaluation

When evaluating the test performance of our model, we computed the precision, recall, and F1 score for each class in each fold. After computing the metrics for each fold, we calculated the mean and standard deviation of each metric across folds. Equations 1, 2, and 3 show the formulas for precision, recall, and F1 score, respectively.

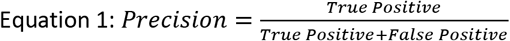

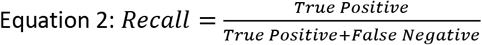

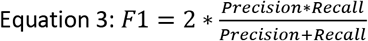

### Explainability – Identifying Clusters of Filters

To better understand the global extraction of features by the model, we selected the model from the fold with the highest weighted F1 score on the test data. We then used a Fast Fourier Transform (FFT) to convert the 30 filters in the first convolutional layer into the frequency domain. Next, we calculated the spectral power between 0 and 50 Hz. We then performed two rounds of k-means clustering using scikit-learn (36). In the first round of clustering, we used 50 initialization and applied the silhouette method to determine the optimal number of clusters. After determining the optimal number of clusters, we redid the clustering with 100 initializations with the optimal number of clusters and examined the spectra of the filters within each cluster.

### Explainability – Examining Importance of Each Cluster of Filters

After identifying clusters of filters, we sought to understand the relative importance of each of the clusters. We applied two methods to this end: ablation and layer-wise relevance propagation (LRP). In our ablation approach, we replaced each cluster of filters with zeros and measured the percent change in the weighted F1 score and F1 scores for each class following ablation. A large negative percent change in performance after ablation corresponds to increase cluster importance.

LRP (6) is a popular gradient-based feature attribution method (37). Rather than examining the effect of perturbing the model, it utilizes the gradients and activations of the network to estimate relevance (i.e., importance). LRP can output both positive and negative relevance. Positive relevance indicates that particular features provide evidence for a sample being assigned to the class it is ultimately assigned to by the classifier. In contrast, negative relevance indicates features that provide evidence for a sample being assigned to classes other than what it is ultimately assigned to by the classifier. In this study, we used the αβ relevance rule with an α of 1 and a β of 0 to filter out all negative relevance and only propagate positive relevance. Equation 4 shows the αβ-rule.

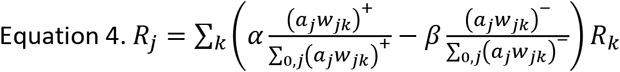

Where the subscript *k* corresponds to a value for one of *K* nodes in a deeper layer and *j* corresponds to a value for one of *J* nodes in a shallower layer. The activation output by the shallower layer is referred to as *a_j_*, and the model weights are referred to by *w*. The relevance is split into positive and negative portions when propagated backwards. The variables α and β control how much positive and negative relevance are propagated backwards, respectively.

LRP typically gives relevance values for individual features, but we wanted to gain insight into filter importance. As such, we computed the relevance for the first layer convolutional layer activations of all test samples, computed the percentage of relevance assigned to each cluster for each subject, summed the relevance values within each predicted class and cluster, and then scaled the relevance values of each respective cluster by the total number of filters minus the number of filters in each cluster divided by the total number of filters. After understanding the relative importance of each cluster of filters, we sought to understand the important components of each cluster of filters. Equation 5 shows how we computed the importance for each class and cluster.

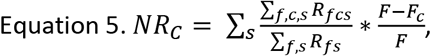

Where *NR_C_* indicates the normalized relevance within a particular cluster (C) and class, *R* indicates relevance, *f* indicates a filter, *c* indicates a cluster, *s* indicates a subject within cluster *C, F* indicates the total number of filters, and *F_C_* indicates the total number of filters in a cluster.

### Explainability – Examining Why Filter Clusters are Important Spectrally

While visualizing the frequency domain of each cluster of filters provided insight into the frequencies learned by the classifier, they did not necessarily indicate the relative importance of each of the frequency bands within each cluster of filters. As such, we performed a separate analysis to understand the relative importance of each frequency band within each cluster of filters. To this end, we iteratively converted each cluster of filters to the frequency domain using an FFT, assigned the coefficients associated with a particular frequency band values of zero, converted the perturbed frequency domain values back to the time domain, and examined the change in class-specific F1 score across the test data. When perturbing the frequency domain of the filters, we perturbed 5 frequency bands – δ (0 – 4 Hz), θ (4 – 8 Hz), α (8 – 12 Hz), β (12 – 25 Hz), and γ (25 – 50 Hz).

### Explainability – Examining Why Filter Clusters are Important Temporally

To understand the relative importance of the waveforms within each of the filters, we used a sliding window approach to iteratively ablate filter weights and calculate the percent change in class-specific F1 score. The sliding window had a window length of 25 points and a step size of 1 point. We assigned the F1 score for each perturbation to the point at the center of each window. To ensure that each individual feature weight had an assigned importance value, we zero-padded the filter prior to the sliding window ablation process.

## Results

In this section, we describe the performance of our model and the results of each of the explainability analyses that we performed.

### Model Performance

Table 2 shows our model performance results. The performance of our model was reasonable overall. The classifier generally had highest performance for the Awake class, followed by performance for NREM2. Additionally, performance for Awake and NREM2 had a low standard deviation and was generally consistent across folds. Performance for NREM3 and was comparable to REM. NREM3 had a noticeably higher mean recall than REM and a slightly higher mean F1 score than REM. REM had a markedly higher mean precision than NREM3. However, REM had much lower levels in variation of performance across folds than NREM3. Interestingly, NREM1 performance was above chance level but still much lower than the performance for all other classes across metrics. NREM1 had lower levels of variation in precision and the F1 score across folds but higher levels of variation in recall.

**Table 2.** Model Test Performance

|  | Awake | NREM1 | NREM2 | NREM3 | REM |
| --- | --- | --- | --- | --- | --- |
| **Precision** | 94.90±02.67 | 38.47±04.85 | 79.83±04.56 | 63.02±12.25 | 67.27±08.53 |
| **Recall** | 89.12±02.44 | 41.10±10.46 | 79.12±05.77 | 75.55±15.04 | 68.65±06.44 |
| **F1** | 91.88±01.82 | 38.56±04.37 | 79.20±01.67 | 67.89±11.09 | 67.41±04.80 |

### Results for Clustering Spectra

We identified 3 clusters with our spectral clustering approach. Figure 3 shows the filters for each cluster in the time domain and in the frequency domain. Clusters 0, 1, and 2 were assigned 3, 16, and 11 of the 30 filters, respectively. Cluster 0 was the smallest cluster. It contained sinusoids of varying amplitude that were predominantly associated with the lower β band, although one of the filters had some α activity. Cluster 1 was the largest cluster. Although it contained a mixture of frequencies, it was predominantly composed of upper β band activity and some upper θ and lower α. In contrast to the other clusters, Cluster 2 was primarily composed of filters extracting δ and lower θ-band activity. Interestingly, several filters did extract lower β-band activity. There were a number of dominant low frequency waveforms in Cluster 2 filters that were not purely sinusoidal (e.g., in filters 20, 21, 26, and 28).

**Figure 3.**
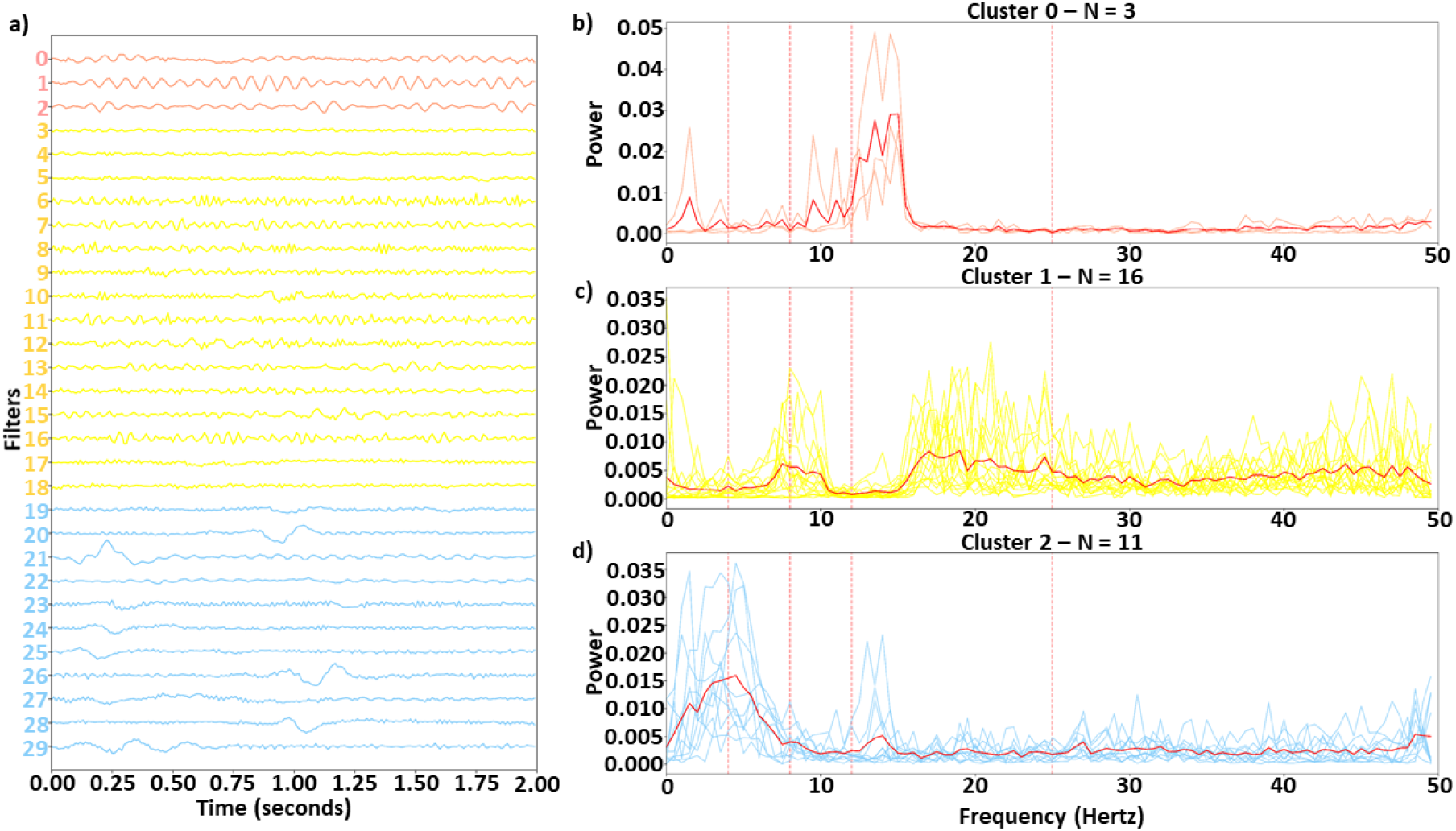
Identified Clusters in Time and Frequency Domain. Panel a shows the time domain of the identified clusters, while panels b, c, and d show the frequency domain of the filters in Clusters 0, 1, and 2, respectively. In panel a, the filters are arranged vertically based on cluster. The solid red lines in panels b through d are the cluster centers, and the vertical dashed lines show boundaries of each frequency band. Light red, yellow, and light blue lines represent clusters 0, 1, and 2, respectively.

### Results for Cluster Importance

Figure 4 shows the overall importance of each cluster to the classifier using both LRP and ablation. With a few exceptions both methods yield similar results. The results are the same for NREM2, NREM3, and REM. They show that Cluster 2 is most important across the three classes. Additionally, across both methods Cluster 1 is second most important for REM and NREM3, while Cluster 0 is second most important for NREM2. For NREM1 and Awake, there are some key differences. Namely, LRP finds Cluster 2 followed by Cluster 1 to be most important. In contrast, ablation finds Cluster 2 followed by Cluster 1 to be most important. For both methods, Cluster 0 is of low to moderate importance for all classes except NREM2. Additionally, the weighted F1 score of the ablation indicates that Clusters 1 and 2, followed by Cluster 0, are most important.

**Figure 4.**
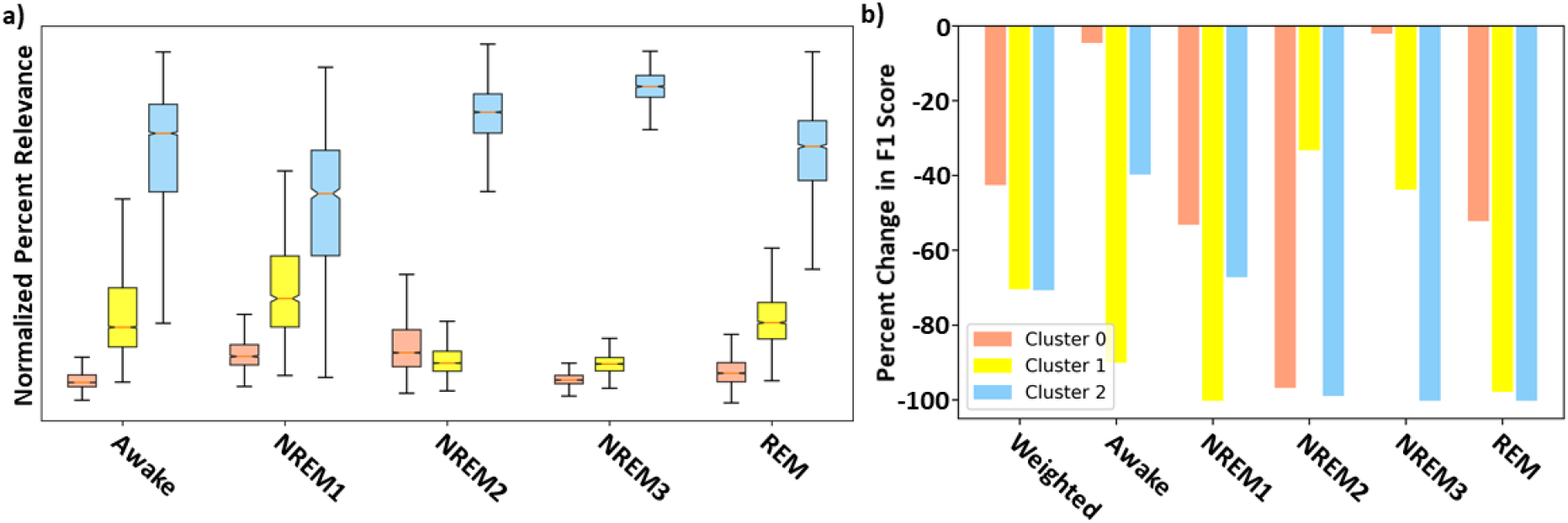
Filter Cluster Importance with LRP and Ablation. Panels a and b show the global importance of each cluster of filters using LRP and ablation, respectively. Importantly, red, yellow, and blue values show importance of Clusters 0, 1, and 2, respectively.

### Results for Cluster-Specific Spectral Perturbation

Our previous analyses characterized and identified the relative importance of each cluster of filters. To examine the importance of different spectra within each cluster, we perturbed the frequency bands in each cluster and examined their effect on classifier performance. Figure 5 shows the results for our spectral perturbation analysis. β was the most important band within Cluster 0. While there were small levels of δ and α activity (as shown in Figure 3) in the cluster, the bands had little to no importance. Similar to in Cluster 0, β was the most important band in Cluster 1, having strong effects on NREM1 and minor effects on NREM3 and REM. Importantly, δ and θ were most important in Cluster 2. Perturbation of δ had strong effects upon all classes except Awake, and perturbation of Θ had a strong effect on REM, with low to moderate effects on NREM1 and NREM2. Interestingly, the Awake class was not strongly affected by the perturbation of frequency bands in any of the classes. The strongest effects of spectral perturbation across clusters and bands were those of Cluster 2 δ and θ upon REM.

**Figure 5.**
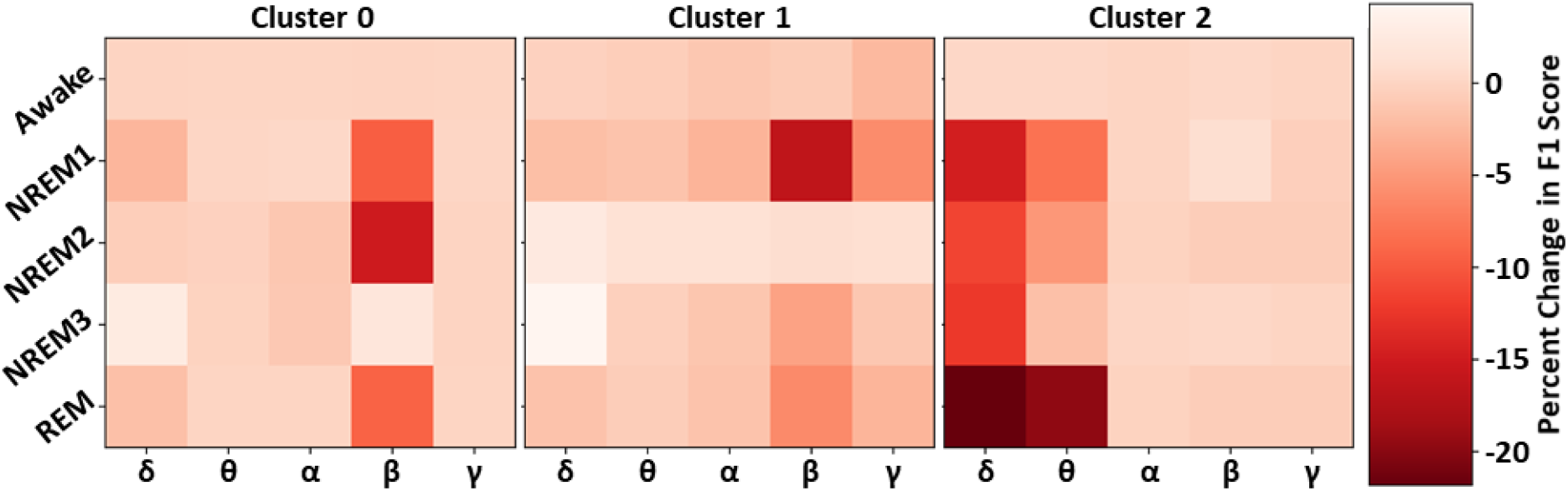
Effect of Cluster-level Spectral Perturbation on Each Class. The leftmost, middle, and rightmost panels show the percent change in class-specific F1 score that resulted from perturbing each frequency band in Clusters 0, 1, and 2, respectively.

### Results for Temporal Filter Ablation

After examining the key frequencies of each cluster of filters, we examined the waveforms extracted by the filters. Figure 6 shows the results of our temporal ablation analysis. The change in weighted F1 score identified the overall impact of the temporal ablation upon the classifier performance. Ablation of Cluster 1 time windows had a slight impact upon classifier performance across most filters and time windows, while ablation of Cluster 0 time windows had no noticeable effect. Cluster 2 had large levels of localized importance in parts of filters 21, 26, and 29.

**Figure 6.**
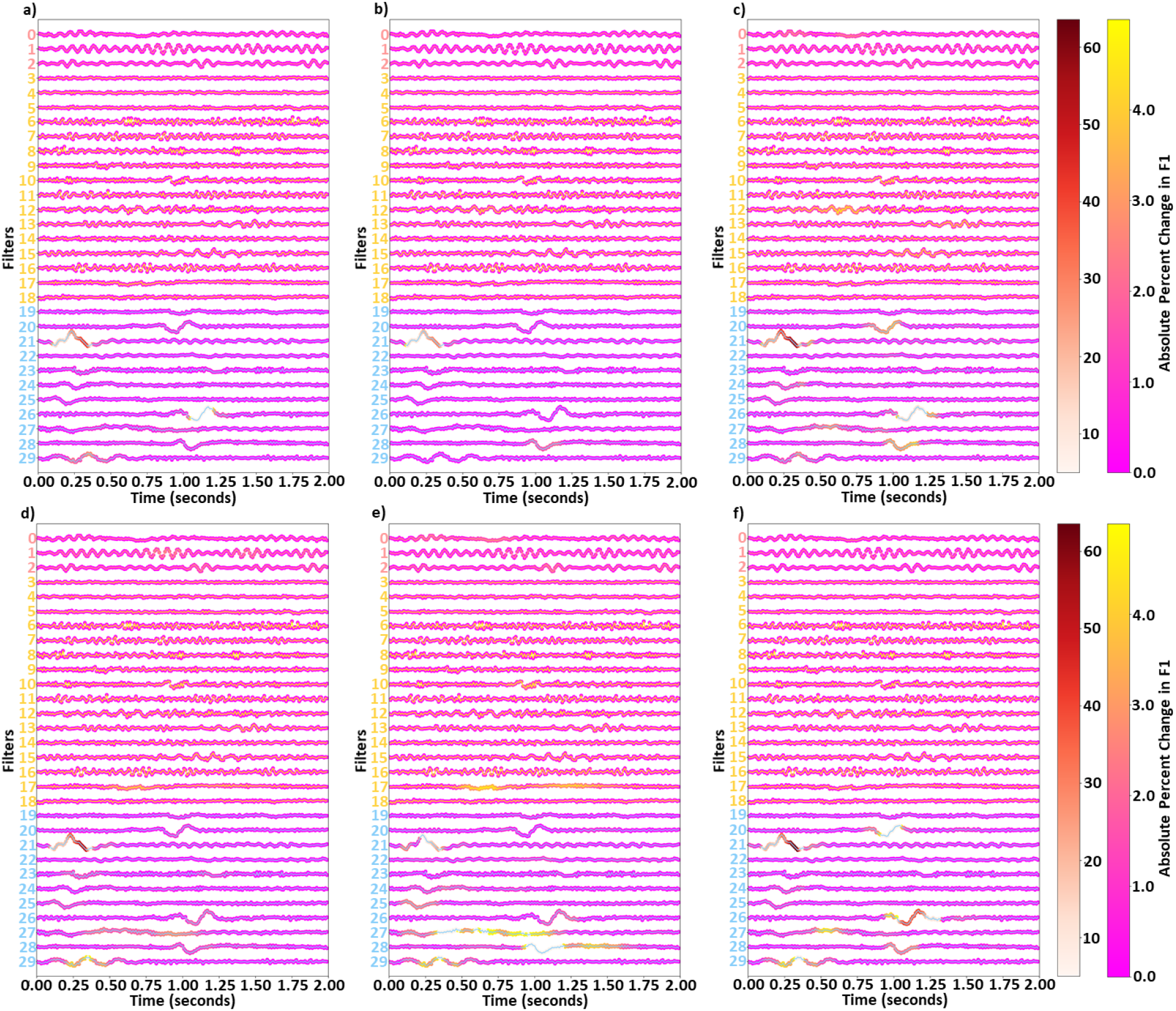
Effects of Filter Temporal Ablation. Panels a through f show the effect of ablation upon the weighted F1-score and Awake, NREM1, NREM2, NREM3, and REM class-specific F1-scores, respectively. Red, yellow, and blue lines show weights for Clusters 0, 1, and 2, respectively. Markers at each time point and the two colormaps indicate the effect of ablating the surrounding 25 time points upon the F1-score. There are 2 colormaps to enable better visualization of importance for both low and high levels of importance.

Similar waveforms showed impacts upon class-specific F1 scores. However, there were also some variations. Interestingly, timepoints 0.8 to 1.1 seconds of filter 20 were of modest importance for NREM1 and REM but not for other classes. Timepoints from 0 to 0.5 seconds of filter 21 in Cluster 2 had high importance across all classes, with highest importance for NREM1, NREM2, and REM and noticeably less importance for Awake and NREM3. Timepoints 1.00 to 1.25 seconds of filter 26 had little importance for Awake but moderate (i.e., NREM1, NREM2, NREM3) to high (i.e., REM) levels of importance for other classes. Timepoints 1.0 to 1.25 seconds of filter 28 were important for NREM1, NREM2, and REM. Timepoints 0.25 to 0.5 seconds of filter 29 had moderate levels of importance for NREM2, NREM3, and REM. Interestingly, timepoints 0.5 to 1.0 seconds of filter 17 were important for NREM2 and NREM3, with other parts of the filter also being important for NREM3. Part of filter 12 were also noticeably important for NREM1.

## Discussion

In this section, we discuss how the results from our analyses relate to one another to explain the model more effectively and how those results compare to existing knowledge from the sleep domain. Unless otherwise specified, when we discuss the results within the context of domain knowledge, we are comparing them to the AASM Manual (28).

### Developing a High Performing Classifier

Classifier performance was highest for the Awake class, which makes sense given that the Awake class had the largest number of samples and that it has clear differences from the other classes. Although NREM1 performance was low, that is acceptable. Most sleep stage classification studies have difficulty classifying NREM1 effectively (38–41), possibly because it is one of the smaller sample groups and can appear similar to Awake and REM (28)(39). Some studies have gone so far as to develop hierarchical models specifically designed to improve performance for NREM1 classification (38). After Awake, the performance of the classifier for NREM2, NREM3, and REM seem to be loosely related to the number of samples in each class.

### Identifying Filter Clusters with Distinct Spectral Features and Quantifying Their Relative Importance

In our first analysis, we visualized the first layer filters of the model from the fold with the highest F1-score. We then clustered the filters in the frequency domain to identify sets of spectrally distinct filters. Interestingly, all canonical frequency bands were present in at least one of the filter clusters. Cluster 0 contained large amounts of lower β activity and is highly important for identifying NREM2. This makes sense given that NREM2 often contains sleep spindles that appear in the lower β band (i.e., 12 – 14 Hz). Cluster 0 was also important for identifying NREM1 and REM and was overall least important for identifying classes other than NREM2. Interestingly, previous studies have shown increased levels of β activity during REM (42). Cluster 1 was characterized as having some lower frequency activity (upper θ and lower α) but predominantly higher frequency activity (upper β and γ). It was very important for identifying Awake, NREM1, and REM samples. The often erratic and a high frequency activity of Awake could explain why the cluster was important for identifying the Awake class. Incidentally, NREM1 is often characterized by low amplitude, mixed frequency activity within the θ range, which could indicate why the cluster was important for NREM1, and as REM is often characterized as multispectral, it makes sense that a cluster having multiple distinct frequency bands would be important for identifying REM. Cluster 2 was characterized by low frequency δ and θ activity and was very important for identifying NREM2, NREM3, and REM, with moderate importance for NREM1. Its importance for NREM1 could be attributed to its extraction of low amplitude θ activity. NREM2 often includes K-complexes that appear within the δ band. Given that Cluster 2 extracts δ activity and extracts waveforms in a number of filters (i.e., 20, 23, 24, 25, 26, and 28) that resemble k-complexes, it is very reasonable that Cluster 2 would be important for identifying NREM2. Importantly, the main feature of NREM3 is δ activity, which could explain why Cluster 2 is so important for NREM3. Lastly, REM has previously been associated with high levels of frontal θ activity (42).

### Confirming Spectral Importance of Clusters

Based on our initial identification of clusters of filters and the relative importance of the clusters for each class, we suggested a number of reasons why the filters might be important to particular classes. However, our previous cluster ablation analysis did not provide definitive evidence regarding the importance of the spectral features present in each cluster. Here, we examine the effects of perturbing the canonical frequency bands within each cluster and examine their relative impact upon classifier performance. Overall, the effect of perturbing the individual bands within each cluster does not sum up to the effect of ablating each cluster, which could indicate that all of the useful information in the filters was not found in the spectral features extracted. For example, the perturbation of frequency bands across all clusters did not affect performance for Awake, which could indicate waveforms, rather than frequency bands, were important for identifying Awake.

Our previous cluster importance analysis indicated that Cluster 0 was highly important for NREM2, with moderate importance for NREM1 and REM, and little to no importance for Awake and NREM3. Our spectral perturbation analysis confirmed that β was the only important frequency band in Cluster 0. Additionally, perturbing β in Cluster 0 had a larger impact upon NREM2 than the perturbation of any other band in Clusters 1 and 2. REM is often characterized as having high β activity (42), which could explain the importance of Cluster 0 β upon REM, but NREM1 is not typically associated with β, which could indicate that the classifier incorrectly associated β with NREM1 and could explain the poor performance of the classifier for the sleep stage. The effect of perturbing β in Cluster 1 upon NREM1 was the largest effect of any pair of classes and bands within Cluster 1. Interestingly, perturbation of individual frequency bands in Cluster 1 seemed to have very little impact upon Awake and REM. Given the importance of the cluster for identifying the stages, that could indicate that the classifier did not rely solely upon extracted spectral features or upon any single frequency band within the cluster when identifying Awake and REM. Instead, the classifier might rely more upon extracted waveforms. The perturbation of δ and θ in Cluster 2 was particularly impactful across multiple classes, particularly the REM class that is characterized as having high levels of θ activity (42). Similar to our previous hypotheses and consistent with clinical guidelines, perturbing θ in Cluster 2 did impact NREM1 performance, and perturbing δ impacted NREM3 performance.

### Examining Importance of Extracted Waveforms

Our analysis of the importance of the canonical frequency bands within each cluster did not fully explain the importance of each cluster to the individual sleep stages. As such, to more fully understand the importance of each cluster, we sought to examine the importance of individual waveforms within the filters to performance for each class. The importance of Cluster 0 to NREM2 was primarily explained by the extraction of β activity. However, by perturbing filters 1 (i.e., 0.75 to 1.25 seconds) and 2 (1.0 to 1.25 seconds) in Cluster 0, we can see some time windows where waveforms resembling sleep spindles seem to be of some importance. While perturbing individual frequency bands in Cluster 1 seemed to have little impact, the perturbation of individual waveforms in Cluster 1 seemed to generally have more of an impact than the perturbation of most time windows in other clusters, which could indicate that the temporal characteristics of the filters were more important than their spectral characteristics. This could also explain the importance of Cluster 1 for identifying Awake and NREM1. Lastly, as Cluster 2 had more low frequency activity, there were more clearly discernable waveforms of strong importance to the classifier performance. Perturbation of filter 21 (i.e., 0.00 to 0.50 seconds) was of high but varying importance across all classes. Low frequency oscillations in a several filters (i.e., 27 and 29) were important for identifying NREM2. Additionally, waveforms resembling k-complexes in multiple filters (i.e., 23, 25, 26, 28) were also of some importance for identifying NREM2. Low frequency activity was present across multiple filters. Like for NREM2, the classifier relied heavily upon δ waveforms when identifying NREM3. Additionally, Filter 21, which has a waveform resembling a vertex sharp wave from (0.00 to 0.50 seconds), was particularly importance across nearly all classes, and was most important in NREM1, NREM2, and REM. The importance of vertex sharp waves in NREM1 could explain its importance for the class.

### Limitations and Next Steps

One of the key contributions of this paper is the combination of a model architecture that is structured to enable increased interpretability with a systematic approach for examining the key spectral and temporal features learned by the classifier. While the filter size of our first layer enabled the visualization of the extracted filters, it also made the training and evaluation of the architecture very computationally intensive. Future iterations of this analysis approach could likely find sufficient levels of explanatory insight with filter lengths equivalent to 0.5 or 1.0 seconds of signal, while also having a model that could be trained and evaluated with more computational efficiency. Additionally, this study only used data from one electrode. The use of one electrode is common in sleep stage classification, but less so in other domains of EEG analysis. Future iterations of this approach could be generalized to multichannel data by perturbing filters when they are applied to individual channels but not to other channels. Lastly, we applied our approach to sleep stage classification so that we could evaluate its efficacy within a well-characterized domain. In the future, our approach might be applied for biomarker identification in domains that are poorly characterized. It would also be possible to examine the effect of perturbation upon the probability of individual samples belonging to a class or upon subject-specific performance metrics for the purpose of personalized biomarker identification.

